# Biological evaluation and molecular docking study of Ferrociphenol as an anti-melanogenic agent

**DOI:** 10.1101/2025.06.20.660698

**Authors:** Emna Ketata, Aissette Baanannou, Pascal Pigeon, Wajdi Ayadi, Aref Neifar, Siden Top, Saber Masmoudi, Mehdi El Arbi, Gérard Jaouen, Ali Gargouri

## Abstract

Cutaneous hyperpigmentation disorders are associated with abnormal accumulation of melanin pigments, which can be treated using depigmenting agents. In the present study, we investigated the effect of ferrociphenol (Fc-diOH), an organometallic intermediate used for the synthesis of hydroxy-ferrocifen derivatives, which has previously been shown as an inhibitor of Sepia tyrosinase activity, on the inhibition of melanogenesis in B16F10 melanoma cells. Cell viability, melanin quantification and tyrosinase activity assay demonstrated that Fc-diOH treatment reduced the amount of intracellular melanin and tyrosinase activity by 32 and by 25%, respectively, in B16F10 melanoma cells at 25 nM without significant cellular toxicity. Furthermore, the biological activity of Fc-diOH against melanogenesis was confirmed in *in vivo* experiments using zebrafish *Danio rerio* embryos. We found that Fc-diOH inhibited melanin production and tyrosinase activity of zebrafish embryos treated with 0.5 and 2 µM respectively, without affecting embryonic development or viability. In addition and interestingly, molecular docking analysis demonstrates that the *p*-hydroxyphenyl groups of Fc-diOH make close contacts with the active site of tyrosinase, compared to arbutin and phenylthiourea, which could be due to its structural homology with the tyrosinase substrate. Therefore, these results strongly suggest that Fc-diOH decreases tyrosinase activity, thereby negatively regulating melanogenesis in B16F10 cells and zebrafish embryos. Thus, Fc-diOH could be used as a depigmentation agent for the treatment of various hyper-pigmentation disorders.

## 1. Introduction

Melanin is a biopolymer pigment synthesized in melanosomes (membrane-bound organelles) within the melanocytes, via a process called melanogenesis. The type, the distribution and the amount of melanin in the melanocytes present in the basal layer of the epidermis and in hair follicles determine the actual color of the skin and hair color of humans and animals [1–3]. Melanogenesis combines enzymatically catalyzed and spontaneous chemical reactions. Tyrosinase catalyzes the rate-limiting step in melanogenesis. It catalyzes the hydroxylation of L-tyrosine to 3, 4 dihydroxy-phenylalanine (L-DOPA) and of L-DOPA to dopaquinone [4]. Following the production of dopaquinone, two main pathways branch off, ultimately resulting in the synthesis of two types of melanin, brown-black eumelanin and yellow red pheomelanin [5, 6]. In the case of eumelanin, Dopaquinone is converted through a redox exchange into L-dopachrome and then to mixtures of 5, 6-dihydroxyindole (DHI) and 5, 6 dihydroxyindole-2-carboxilic acid (DHICA) mixtures, according to the decarboxylation rate and Trp2 activity respectively. Finally, a new oxidation of these dihydroxy-indoles by tyrosinase or Trp1 gives rise to indole quinones which polymerize and give rise to the eumelanin.

Functionally, melanin protects the skin from various types of ionizing radiation, including ultraviolet (UV), and plays a crucial role in the absorption of free radicals [7]. However, genetic, hormonal and environmental factors can lead to an excessive production of melanin in the skin [8–10] and causes diseases including melanoma and pigmentary disorders, such as melasma, age spots, and sites of actinic lesions [11].

Therefore, several melanogenesis inhibitors are currently used as pharmaceutical or cosmetic additives however, due the low stability of their formulation and undesirable side effects (cytotoxicity, skin irritation, carcinogenicity), their use is still limited [12]. Due to these safety concerns, the ongoing search for potential new molecules preventing pigment disruption continues to progress and attracts increasing interest. Various strategies are used to search for inhibitors of melanogenesis and include *in silico*, *in vitro*, and *in vivo* approaches [13–16]. Recently, zebrafish has emerged as a valuable *in vivo* model for evaluating the depigmenting activity of small molecules [17], for screening compounds that controlling pigment cell development [18]. In addition to its numerous advantages, including its small size, abundant offspring, and progeny and high efficiency of drug penetration through the skin and gills, the zebrafish embryo has a great physiological and genetic similarity with the mammals. More importantly, the effect of the chemical compound on melanogenesis can be assessed by simple observation of the formation of pigmentation on the surface of this animal without any complicated experimental procedures [18, 19]. Like other vertebrates, zebrafish have pigment cells from two distinct embryonic sources. Those of the epidermis and dermis arise from the neural crest, while those that make up the outermost layer of the retina, the retinal pigment epithelium, derive from the optic cupula and begin in the retinal epithelium and in the melanophores. Pigment cells grow rapidly and within hours, they become an important feature of the embryo [20]. At the embryonic stage, melanocytes that arise from the neural crest pattern the zebrafish embryo by giving rise to four bands: the dorsal larval stripe, the lateral larval stripe, the ventral larval stripe and the yolk sac stripe. Microphthalmia-associated transcription factor (MITF) appears to be the master regulator of melanocyte identity that controls the melanocytes development from the neural crest [21]. In fact, mice and fish lacking MITF and MITF-a (the fish ortholog of MITF) cannot form melanocytes [22, 23] and ectopic expression of mitf-a can generate ectopic melanocytes [23]. Additionally, MITF has been reported to regulate melanogenesis in the melanocyte lineage by modulating transcription of several genes involved in melanin synthesis including tyrosinase, TRP1 and TRP2 [24], and it appears to be central of a regulatory network of transcription factors and signaling pathways that control the survival, proliferation and differentiation of melanocyte [24].

Fc-diOH, also called ferrociphenol, is an organometallic compound that first served as a simple intermediate for the synthesis of hydroxy-ferrocifen [25], a compound analogous to hydroxy-tamoxifen, which is the bioactive form of tamoxifen, a selective estrogen receptor modulator (SERM) [26]. Although not a SERM itself, due to the absence of the dimethyl-amino-alkyl chain, Fc-diOH has been shown to have a surprisingly strong anti-proliferative effect on human breast cancer cell lines MCF-7 (estrogen receptor positive; ER+) and MDA-MB-231 (estrogen receptor negative; ER-) with an IC_50_ of 0.7 and 0.6 µM, respectively [27]. In fact, almost all compounds having the [ferrocenyl-double bond-*p*-phenol] motif have cytotoxicity activities on various cell lines, but the ferrocenyl and phenol moieties must be in *trans* [28]. For this reason, having two *p*-hydroxyphenyl groups, Fc-diOH still possesses this pharmacophore moiety, as the single *Z* isomer of ferrocifen already does. Some diphenol derivatives have been improved such as the cyclic version (ferrocenophane) of Fc-diOH [28, 29] or a patented [30] succinimide derivative [31], which have IC_50_ below 0.1 µM. For this reason, all compounds bearing this motif, including the ferrocifen itself, have been classified as belonging to a series called the “Ferrociphenol series”. The series leader, Fc-diOH has also been reported to have other biological activities including antimicrobial agent against many species such as *Pseudomonas aeruginosa*, *Staphylococcus aureus*, *Candida albicans* [32], *Pseudomonas savastanoi, Fusarium solani and Aspergillus tumefaciens* [33]. In addition, we recently reported the inhibitory effect of Fc-diOH on, the key enzyme of melanogenesis, Sepia tyrosinase activity; with a competitive inhibition mode and this is due to the structure homology between the substrate (L-tyrosine) and its hydroxy-phenyl group [34]. To our knowledge, no previous studies have reported the effect of Fc-diOH on melanogenesis.

The aim of our study is to investigate the depigmenting effect of Fc-diOH *in vitro* on murine melanoma cell line B16F10 and *in vivo* on zebrafish model. We support this depigmenting effect by a molecular docking analysis between Fc-diOH and tyrosinase.

## 2. Materials and methods

### 2.1. Reagents

L-DOPA, Gentamicin and MTT (Thiazolyl Blue Tetrazolium Bromide) were purchased from BioBasic, Canada. IBMX, arbutin and melanin were obtained from Sigma-Aldrich, Germany. RPMI-1640 Medium, Fetal Bovine Serum and trypsin-EDTA (0.05%) were purchased from Gibco Life Technologies, Paisley, UK. Ferrociphenol (Figure 1) was synthesized as described in [25].

**Figure 1:**
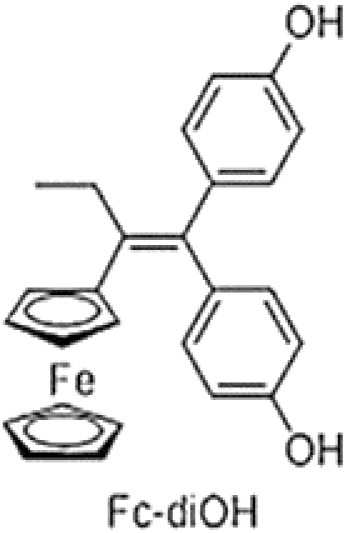
Structure of Fc-diOH.

### 2.2. Murine melanoma cell line culture

The B16F10 melanoma cell line was maintained in RPMI 1640 medium supplemented with 10% fetal bovine serum and gentamicin (50 µg/mL). The cells were cultured in a humidified atmosphere containing 5% CO_2_ at 37°C.

### 2.3. MTT assay

Cell viability was determined using the MTT assay as previously described [35]. B16F10 cells were seeded at a density of 10^4^ cells/well in 96-well tissue culture plate. After 24 h of incubation, the cells were treated with different concentrations of Fc-diOH for 24 h. After media removal, fresh medium containing MTT solution (1 mg/mL) was added and incubation was continued for 4 h at 37°C. Subsequently, the formed formazan crystals formed were dissolved in SDS (10%) and further incubated at room temperature overnight. Absorbance was measured at 570 nm using a microplate reader (Thermo Scientific Varioscan Lux). Data were representative of three independent experiments.

### 2.4. Determination of melanin content

Melanin content was assessed using a previously described method with slight modifications [36]. Briefly, cells were seeded at a density of 2 x 10^5^ in a 6-well plate. After 24 h of incubation, the murine B16F10 cells were incubated without or with Fc-diOH at a concentration ranging from 5 to 25 nM, in the presence of IBMX (3-isobutyl-1-methylxanthin) at 100 µM to induce melanogenesis. After 48 h of incubation, B16F10 cells were harvested and washed twice with PBS. The pelleted cells were dissolved in 1 N NaOH for 1h at 60°C. The mixture was then vortexed vigorously to solubilize the melanin pigment. Then, the absorbance at 410 nm was measured and the melanin content was calculated relative to a known standard of synthetic melanin.

### 2.5. Measurements of intracellular tyrosinase activity

The intracellular tyrosinase activity of B16F10 cells was determined by measuring the oxidation rate of L-DOPA substrate, as previously described [36]. Cells were seeded at a density of 2 x 10^5^ in a 6-well plate. After 24 h incubation, the cells were treated with different concentrations of Fc-diOH, with and without 100 µM IBMX for another 24 h incubation. Cell pellets were washed with PBS, frozen at -20°C and thawed in 200 µl of lysis buffer (10 mM sodium phosphate pH 6.8, 1% Triton X-100, and 1 mM PMSF). Lysates were centrifuged at 13,000 rpm for 20 min at 4°C and the supernatant was used to determine crude tyrosinase activity. Protein content in the supernatant was determined by Bradford assay using BSA as a standard. Fifty micrograms of protein were placed in each well of a 96-well plate and the enzyme assay was initiated by the addition of L-Dopa and sodium phosphate buffer pH 6.8, the absorbance at 475 nm was measured every 10 min for at least 1 h at 37°C using a colorimetric microplate reader. The tyrosinase activity was measured and then compared to the activity of the control induced by IBMX.

### 2.6. Acute toxicity test on zebrafish embryos

Adult zebrafish were maintained in aquaria with an alternating 14/10 hour light/dark cycle at 28°C. Synchronized embryos were obtained from natural reproduction, induced in the morning by turning on the light. Zebrafish embryos were distributed in 24-well plates with 20 embryos per well filled with 2 ml E3 medium and treated with Fc-diOH from 4 h to 48 hpf (hours post-fertilization) with various concentrations ranging from 0 to 10 µM. The observation of sub-lethal and lethal morphological parameters (embryonic coagulation, absence of somite formation, non-detachment of the tail bud, and absence of heart rate) was carried out at 1 and 2 dpf using the Stemi 2000-C stereoscope (Carl Zeiss, Germany) equipped with AxioCam 105 Colors CCD camera (Carl Zeiss).

### 2.7. Evaluation of depigmenting activity in zebrafish embryos

The evaluation of the depigmentation effect in the zebrafish model system was carried out according to the method of Choi [17]. Thirty zebrafish embryos were treated with with 9–48 hpf Fc-diOH at various concentrations (0.1 - 2 µM) and the positive control was treated with 50 µM PTU (*N*-Phenylthiourea). Effects on zebrafish pigmentation were observed under a stereomicroscope (Zeiss Stemi 2000-C) and photographed under a camera (zeiss Axiocam 105 color). Each group was sonicated for 5 min at 70% amplitude in cold lysis buffer (10 mM sodium phosphate pH 6.8, 1% Triton X-100, 1 mM PMSF). The lysate was centrifuged at 12,000 rpm for 20 min at 4°C and the pellet was suspended in 100 μl of NaOH (1N) and incubated at 100 °C for 1 h. The mixture was then vortexed vigorously to solubilize the melanin pigment. The optical density of the supernatant was measured at 410 nm by a colorimetric microplate reader (Thermo Scientific Varioscan Lux).

For the determination of tyrosinase activity, 250 μg of total lysate protein was added to a reaction mixture containing 50 mM phosphate buffer (pH 6.8) and 2.5 mM L-Dopa. After incubation at 37 °C for 60 min, the formation of dopachrome was measured at 475 nm. Tyrosinase activity and melanin content were expressed as the percentage change from that of the control group, which was taken as 100%.

### 2.8. Homology modelling and docking studies on human tyrosinase

The 3D crystal structure of human tyrosinase is not available and the homology modeling was performed to predict the 3D structure of human tyrosinase or hTYR. The amino acid sequence in FASTA format (529 AA, ID: P14679) was retrieved from UniProt Knowledge database (http://www.uniprot.org/uniprot/P14679). As a template tyrosinase from *Bacillus megaterium* (pdb: 3ntm) was chosen. This enzyme has 32.8% sequence identity with the query protein sequence. Homology modeling was carried out on SWISS-MODEL server (SWISS-MODEL (expasy.org). Then, the interaction models of Human tyrosinase with Fc-diOH, arbutin and PTU were predicted by Autodock/Vina [44]. The grid for the ligand conformational search calculations was placed with its center located at the tyrosinase binding site. The docking grid size was 30, 30 and 40 grid points, while the grid centers were designated at dimensions 11, -11 and 6. The best conformations with the lowest binding free energy were selected and analyzed using Discovery Studio 2017 R2 Client.

### 2.9. Statistical analysis

Differences between the results of the melanin content assay and the intracellular tyrosinase activity assay were assessed for statistical significance using the Student’s t-test using SPSS 13.0 statistical software (SPSS). Differences were considered statistically significant at * p<0.05 and ** p<0.01.

## 3. Results

### 3.1. Inhibition of melanogenesis by Fc-diOH in B16F10 cells

The effect of Fc-diOH (Figure 1) on cell viability was evaluated using the MTT assay. The results showed that it had no significant cytotoxic effect on B16F10 cell lines with a concentration range of 5 to 75 nM (Figure 2). A marked reduction in cell viability was observed when the concentration of Fc-diOH was increased to 100 nM. This cytotoxic effect on B16F10 melanoma cells is in agreement with the results previously published by Bruyère et al. [38].

**Figure 2:**
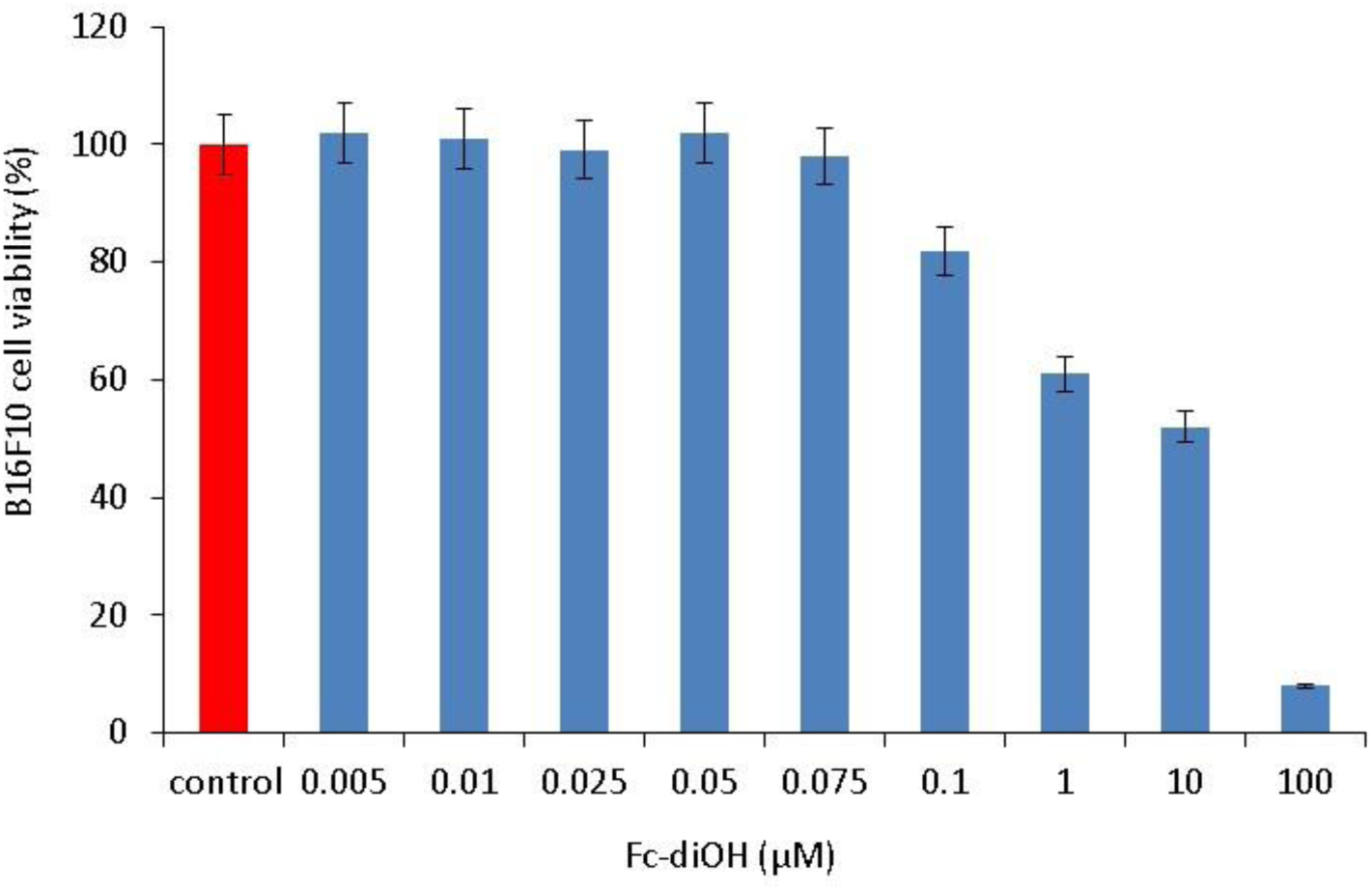
Effect of Fc-diOH on B16F10 cells viability at various concentrations (from 0.005 to 100 µM).

To examine the effects of Fc-diOH on melanogenesis, we determined the melanin contents in B16F10 cells, after induction of melanogenesis by IBMX, followed by the treatment of the cells with Fc-diOH at non-cytotoxic concentrations (5, 10 and 25 nm) and using arbutin as a positive control. Fc-diOH showed a significant inhibitory effect on melanin production in a dose dependent manner (Figure 3.A). Treatment with 25 nM Fc-diOH reduced melanin content in B16F10 cells by 32%, while arbutin treatment used at a much higher dose of 100 µM, induced a slightly stronger effect and reduced melanin production in B16F10 melanoma cells by 36%.

**Figure 3:**
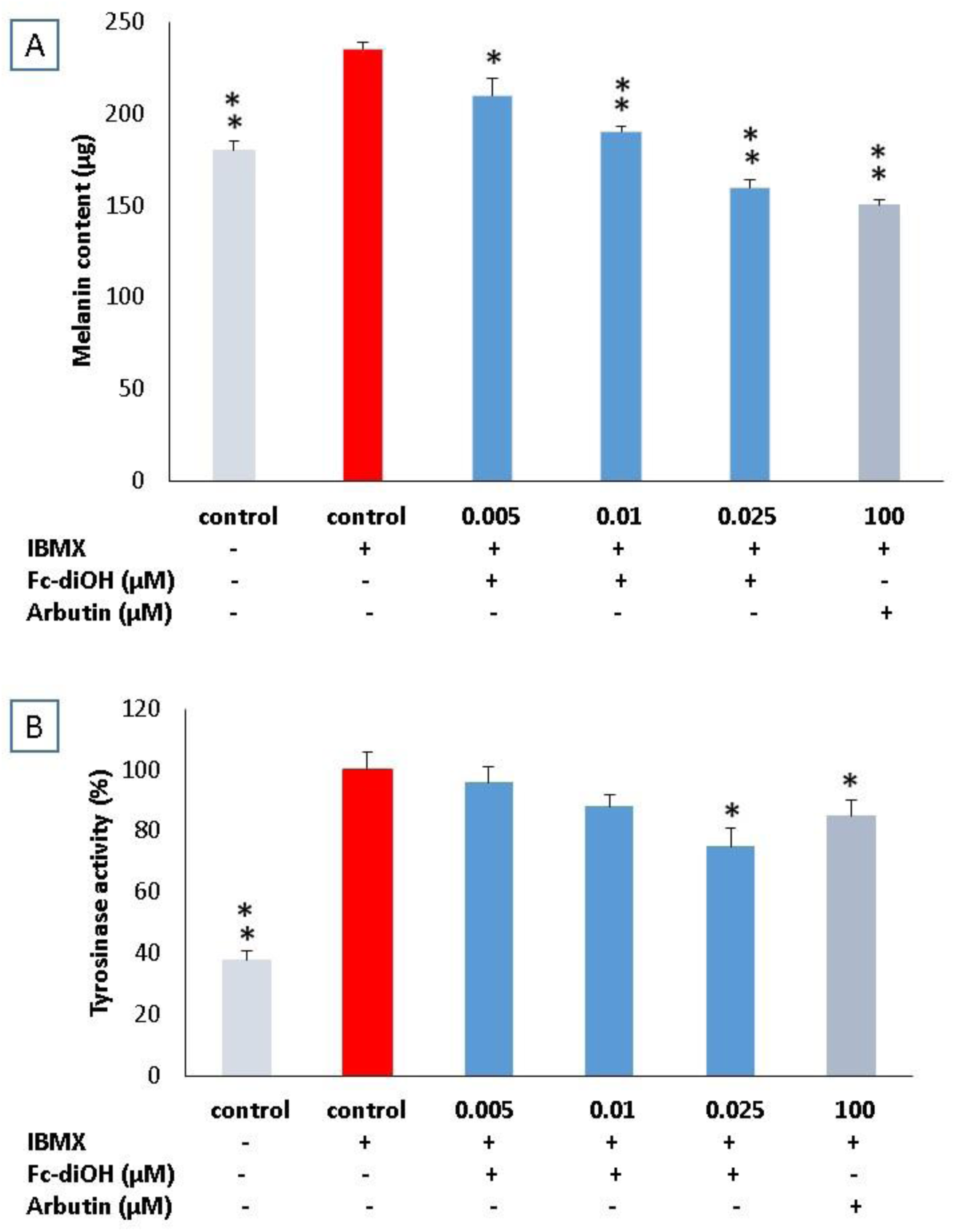
Effect of Fc-diOH on the production of melanin in treated B16f10 cells with 0.005, 0.01 and 0.025 µM concentrations of Fc-diOH and 100 µM of arbutin (A) and the percentages of intracellular tyrosinase activity in B16F10 cells treated with 0.005, 0.01 and 0.025 µM concentrations of Fc-diOH and 100 µM of arbutin (B). Data shown represent the mean ± S.D of three separate experiments. Values are significantly different in comparison with IBMX-treated control. * p < 0.05 and **p < 0.01.

### 3.2. Tyrosinase inhibition in B16F10 cells by Fc-diOH

To assess the effect of Fc-diOH on tyrosinase activity and understand the mechanism by which melanogenesis is inhibited in IBMX stimulated B16F10 melanoma cells, we determined the intracellular tyrosinase activity in B16F10 cells treated with Fc-diOH at 5, 10 and 25 nM and with 100 µM of arbutin as positive control. The results showed that tyrosinase activity was reduced by almost 25% in B16F10 cells treated with 25 nM Fc-diOH (Figure 3.B) and by 15% in cells treated with 100 µM arbutin. Fc-diOH reduced intracellular tyrosinase activity more potently than arbutin, with a lower concentration.

The decrease in tyrosinase activity obtained at 25 nM correlates with the decrease in the amount of melanin described above at this concentration.

### 3.3. Evaluation of the *in vivo* depigmenting effect of Fc-diOH on zebrafish embryos

The zebrafish model was used as an *in vivo* system to evaluated the inhibition of melanogenesis by Fc-diOH (Figure 4). PTU (1-phenyl-2-thiourea) was used as a positive control to inhibit melanin production in zebrafish according to [39] by blocking all steps of tyrosinase-dependent melanin synthesis [20].

**Figure 4:**
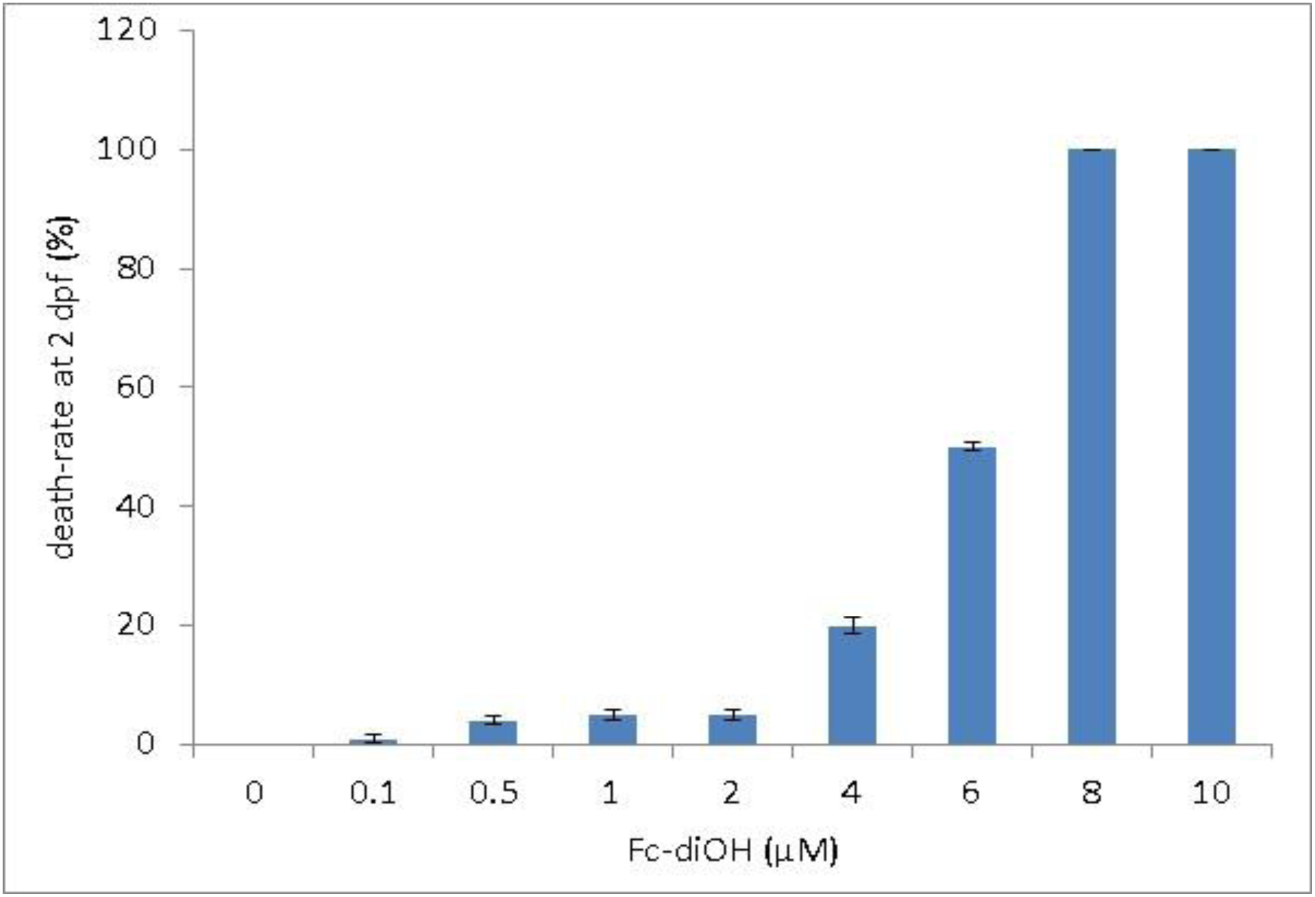
Effects of Fc-diOH on death rate of zebrafish embryos after 2 days post-fertilization (dpf) at various concentrations (From 0.1 to 10 µM).

In order to identify the toxic concentration of Fc-diOH on zebrafish embryos, we performed an acute toxicity test starting from 2 h post fertilization to 2 days post fertilization (Figure 4). Treatment with 1 and 2 µM showed no effect on embryo development and viability. However, treatment with 4, 6 and 8 µM caused lethality of 20%, 54% and 100% respectively.

As shown in Figure 5.A, zebrafish embryos at 1-day post-fertilization (dpf) treated with Fc-diOH had normal development at concentrations of 1 and 2 µM whereas development delay was observed with 6 and 8 µM. Interestingly, treatment with 0.5 and 2 µM of Fc-diOH led to remarkable depigmentation of zebrafish embryos (Figure 5.B). To estimate Fc-diOH inhibitory activities, we measured tyrosinase activity and total melanin content in whole extracts of treated zebrafish embryos. Fc-diOH showed a dose-dependent inhibitory effect on melanin production (Figure 6.A) and tyrosinase activity (Figure 6.B). Fc-diOH induces a depigmenting effect from the concentration of 0.1 µM, well below the effective concentration of PTU necessary for depigmentation.

**Figure 5:**
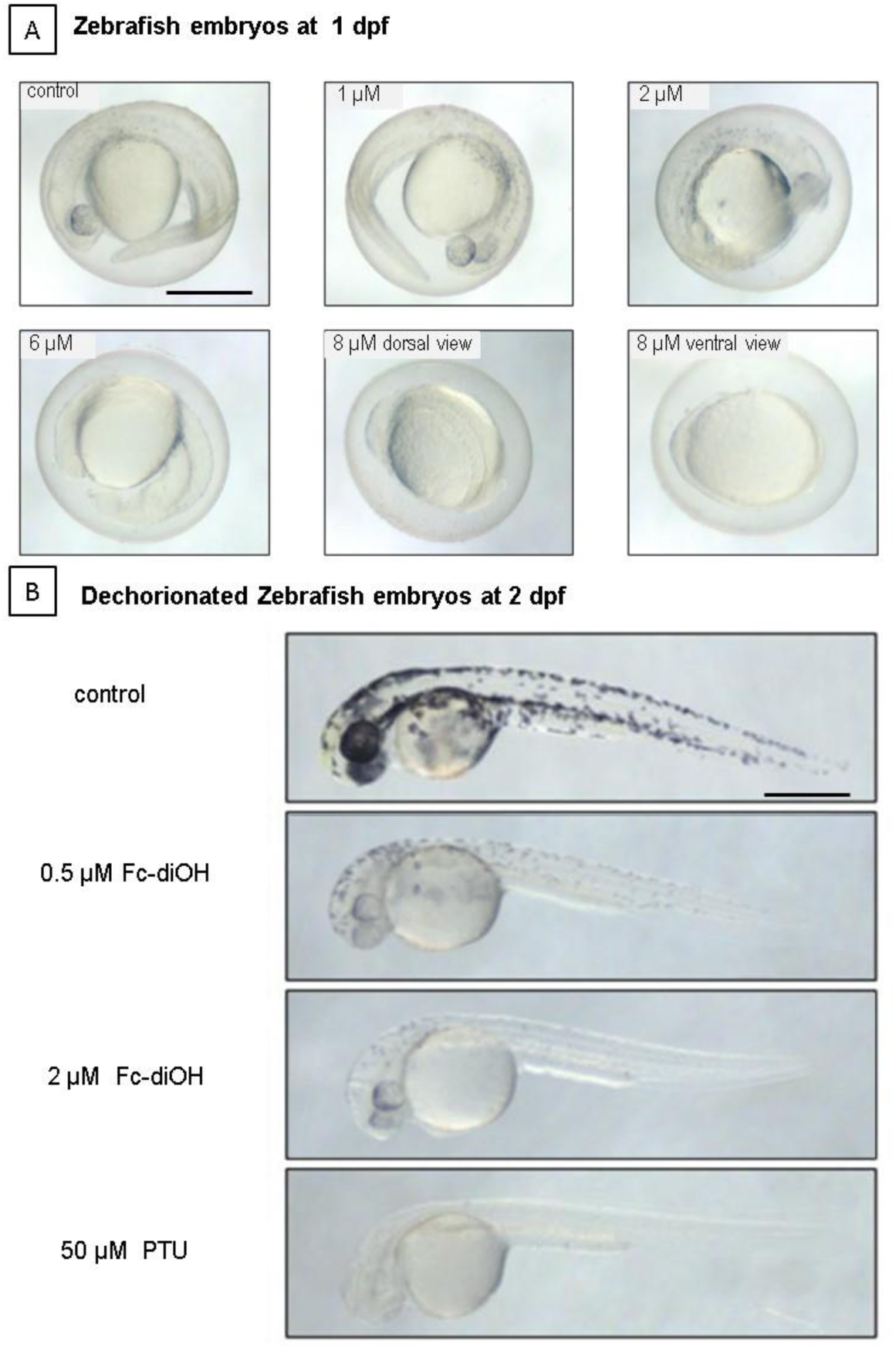
Effects of Fc-diOH on melanogenesis in zebrafish embryos. The effects on the pigmentations of zebrafish embryos were observed by microscopy. (a) zebrafish embryos at 1day post-fertilization (dpf). Untreated embryo: control, treated embryos with 1, 2, 6 and 8 µM of Fc-diOH (b) Dechorionated zebrafish embryos at 2 dpf. Untreated embryo: control, treated embryos with 0.5 and 2 µM of Fc-diOH and treated embryos with 50 µM of 1-phenyl-2-thiourea (PTU).

**Figure 6:**
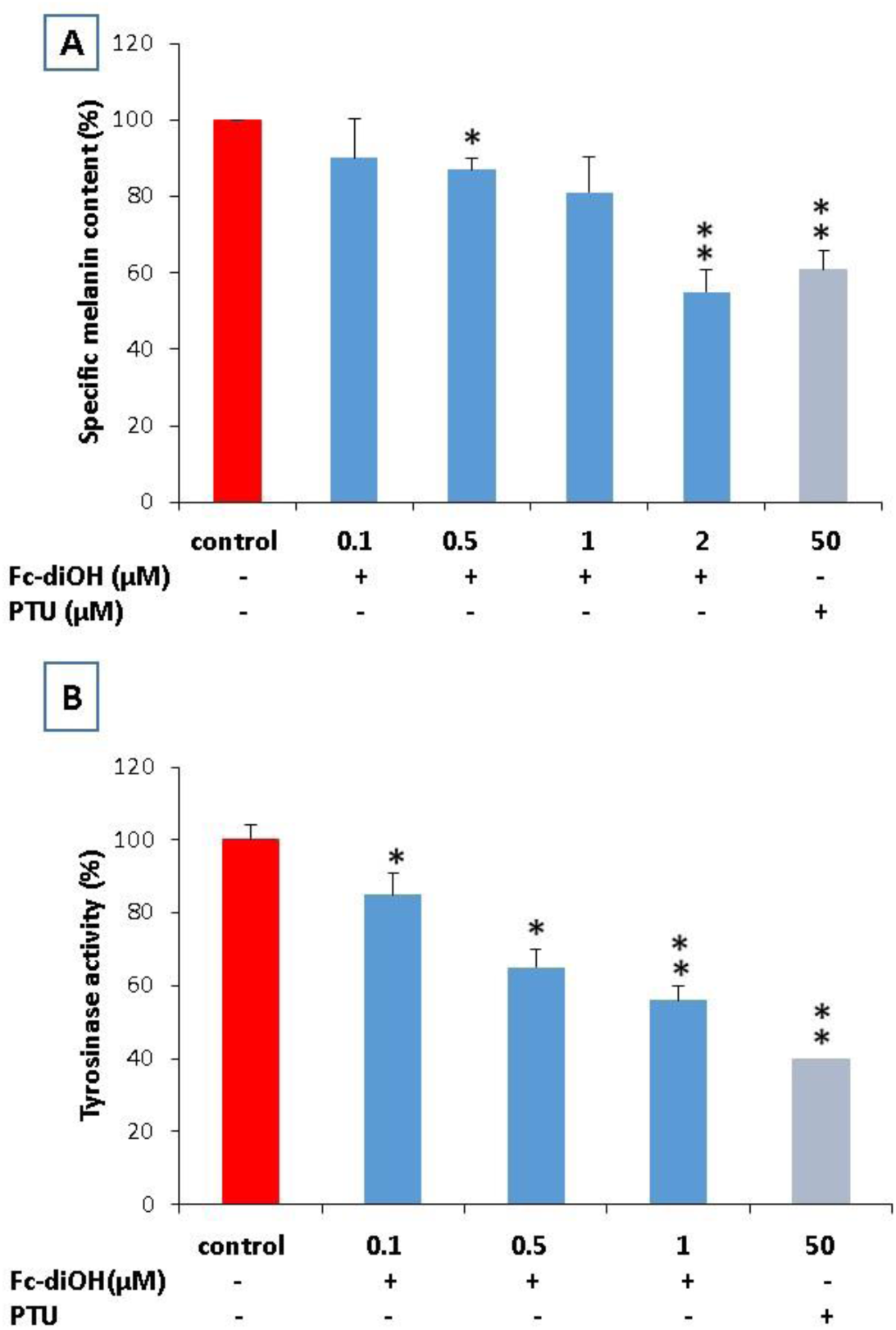
Inhibitory effect of Fc-diOH on the production of melanin in zebrafish embryos at various concentrations from 0.1 to 2 µM of Fc-diOH and 50 µM of PTU (A) and photograph of tyrosinase reaction medium and the corresponding percentage of intracellular tyrosinase activity at various concentrations from 0.1 to 1 µM of Fc-diOH and 50 µM of PTU (B). Data shown represent the mean ± S. D of three separate experiments. Values are significantly different in comparison with the control. * p < 0.05 and **p < 0.01.

### 3.4. Molecular docking analysis

Molecular docking analysis was applied to study the binding interactions of Fc-diOH, arbutin and PTU with the crystal structure of *Agaricus bisporus* tyrosinase or AbTYR (PDB Code: 2Y9X) and with predicted 3D structure of human tyrosinase or hTYR via autodock vina to elucidate the molecular mechanism possible. The catalytic residues of AbTYR were identified as His 85, His 244, Val 248, Asn 260, His 263, Phe 264, Val 283, and His 296 [40]. We show that the p-hydroxyphenyl groups of Fc-diOH interact with the tyrosinase catalytic residues His 263, His 259 and Ser 282 via H-bond interactions and that Fc-diOH interacts with tyrosinase at catalytic residues Val 283, Val 248, Phe 264 via hydrophobic interactions (S. Figure 1. A, B, C and table 1). Additionally, we show that arbutin interacts with tyrosinase at the active site of tyrosinase at residues His 262, Ser 282, His 244 via H-bond interactions (S. Figure 1. D, E, F and Table 1). PTU interacts with tyrosinase at its active site at the catalytic residues His 259, His 244 via H-bond interactions, Val 283 via hydrophobic interaction and His 244 via Pi-Sulfur interaction. Note that the highest binding energy value was for Fc-diOH as -6.6 kcal/mol (Table 1).

**Table 1:** Binding energy and interacting residues of *Agaricus bisporus* tyrosinase (2Y9X) and Human tyrosinase with inhibitors from AutoDock-Vina.

| Compounds | Interactions with *Agaricus bisporus* tyrosinase model (2Y9X) |  | Interactions with Human tyrosinase model |  |
| --- | --- | --- | --- | --- |
|  | Binding energy (KJ/mol) | Interacting residues | Binding energy (KJ/mol) | Interacting residues |
| **Fc-diOH** | -6.6 | **H-bonds:**<br>Ser 282, His 263, His 259<br>**Hydrophobic interactions:**<br>Val 283(2), Val 248, Phe 264 | -8.1 | **H-bonds:**<br>His 304<br>**Hydrophobic interactions:**<br>Ile 368, Phe 347<br>**Electrostatic interaction :**<br>Arg 308 |
| **Arbutin** | -6.2 | **H-bonds:**<br>His 262, Ser 282, His 244 | -6.4 | **H-bonds:**<br>Glu 203, Ser 360<br>**Hydrophobic interactions:**<br>His 202, His 367, Val 377 |
| **PTU** | -4.9 | **H-bonds:**<br>His 259, His 244<br>**Hydrophobic interactions:**<br>Val 283<br>**Pi-Sulfur interactions:**<br>His 244 | -4.5 | **H-bonds:**<br>Gln 286<br>**Hydrophobic interactions:**<br>Gln 376<br>**Pi-Sulfur interactions:**<br>His 285, Cys 289 |

In the second step of our docking study, we sought to predict the 3D crystal structure of human tyrosinase using SWISS-MODEL server (SWISS-MODEL (expasy.org) and using tyrosinase from *Bacillus megaterium or* bmTYR (pdb: 3ntm) as a template (S. Figure 2). The superposition of the hTYR model on bmTYR shared a similar catalytic cavity with substitution of the catalytic residues of bmTYR His 42, His 60, His 69, His 204, His 208, His 231, Val 2018 and Arg 209 [41] by catalytic residues of hTYR predicted model His 180, His 202, His 211, His 363 His 367, His 390, Val 377 and Ile 368 (S. Figure 3). The obtained hTYR model was used to perform the molecular docking studies of Fc-diOH, arbutin and PTU. We show that the p-hydroxyphenyl group of Fc-diOH interacts with the catalytic tyrosinase residues Ile 368 which corresponds to the catalytic residue Arg 209 of bmTYR which play a role in substrate binding orientation, based on their flexibility and position [41] and with residues His 304 and Arg 308 (Figure 7 .A, B, C and table 1). Additionally we show that the hydroxyphenyl group of arbutin interacts with the hTYR catalytic residues His 202, His 367 and Val 377 via hydrophobic interactions (Figure 7. D, E, F and table 1).

**Figure 7:**
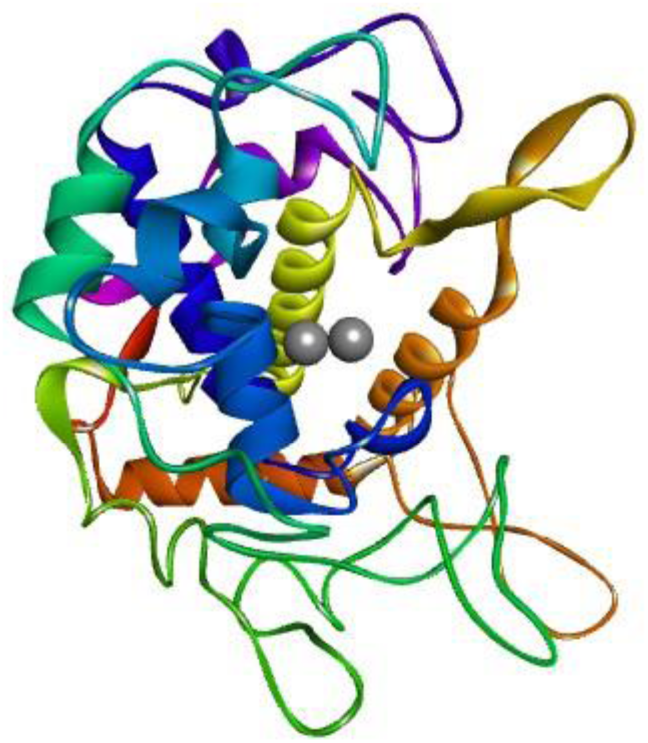
Predicted human tyrosinase model.

**Figure 8:**
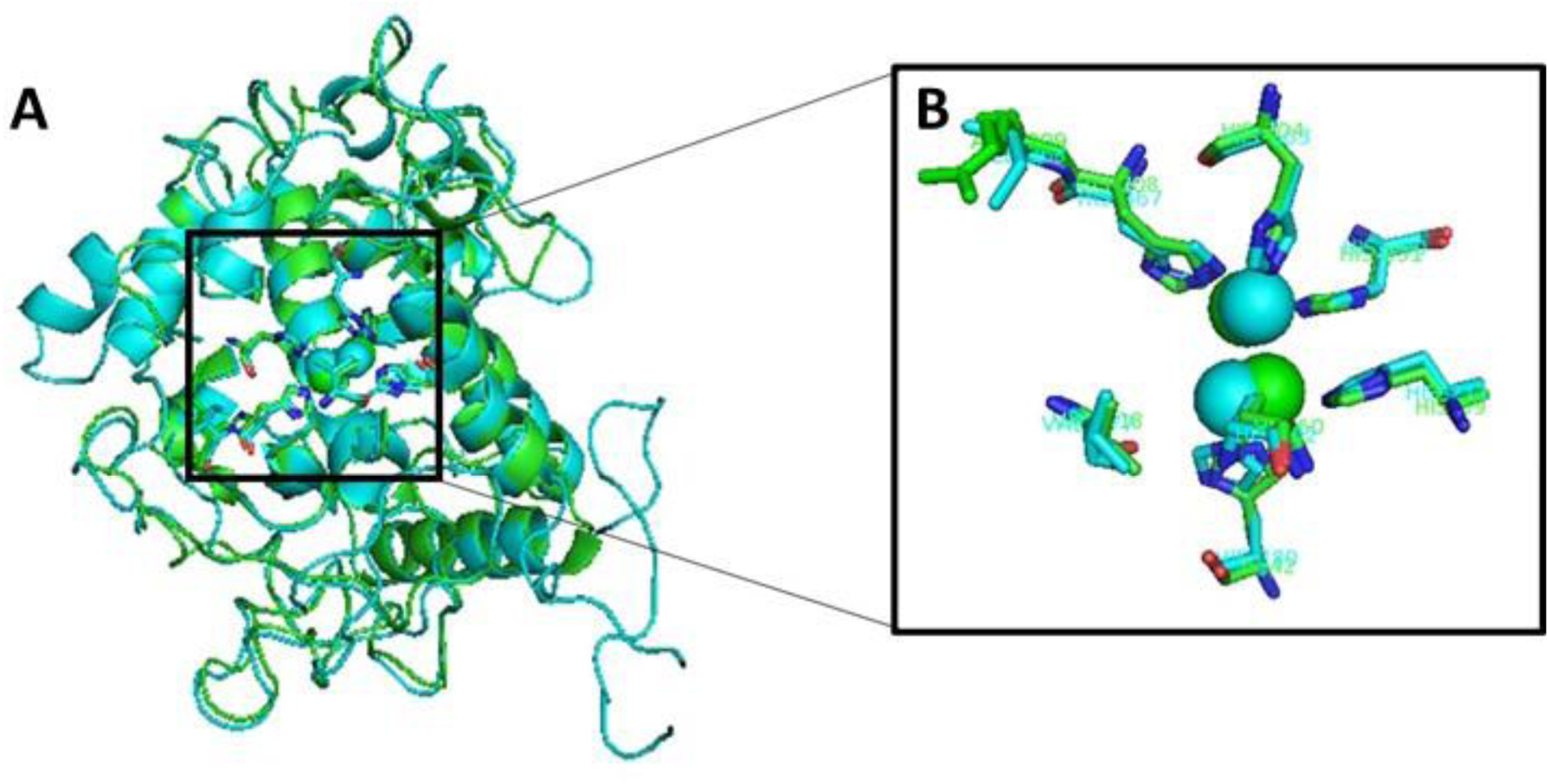
A) Superposition of the hTYR homology model (blue sticks) on bmTYR (green sticks). B) Close view of hTYR and bmTYR active site, the catalytic residues of bmTYR are (His 42, His 60, His 69, His 204, His 208, His 231, Val 2018 and Arg 209) and catalytic residues of hTYR predicted model are (His 180, His 202, His 211, His 363 His 367, His 390, Val 377 and Ile 368).

**Figure 9:**
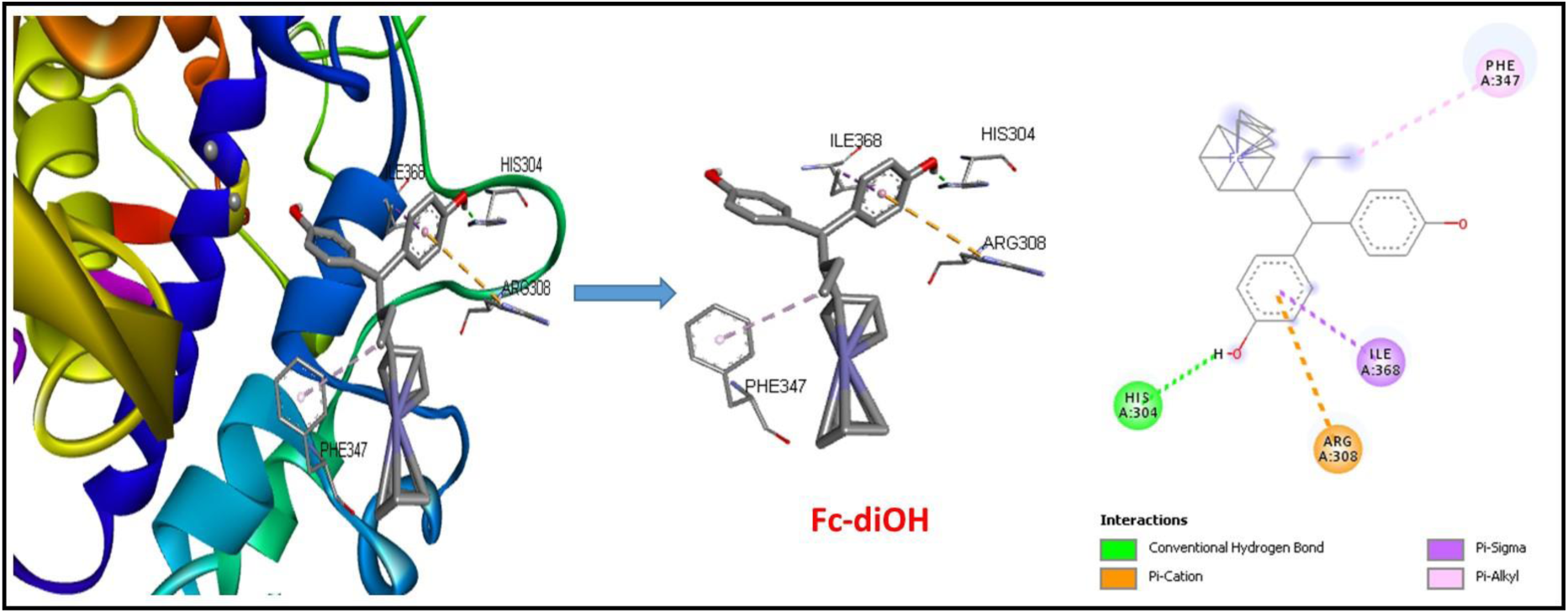
Representation of the putative binding modes of Fc-diOH in the “pocket” of human tyrosinase model. The green dotted lines indicate H-bond interaction, the pink and purple dotted lines indicate hydrophobic interactions, orange dotted lines indicate electrostatic interactions and the two gray spheres indicate copper ions at the active site.

**Figure 10:**
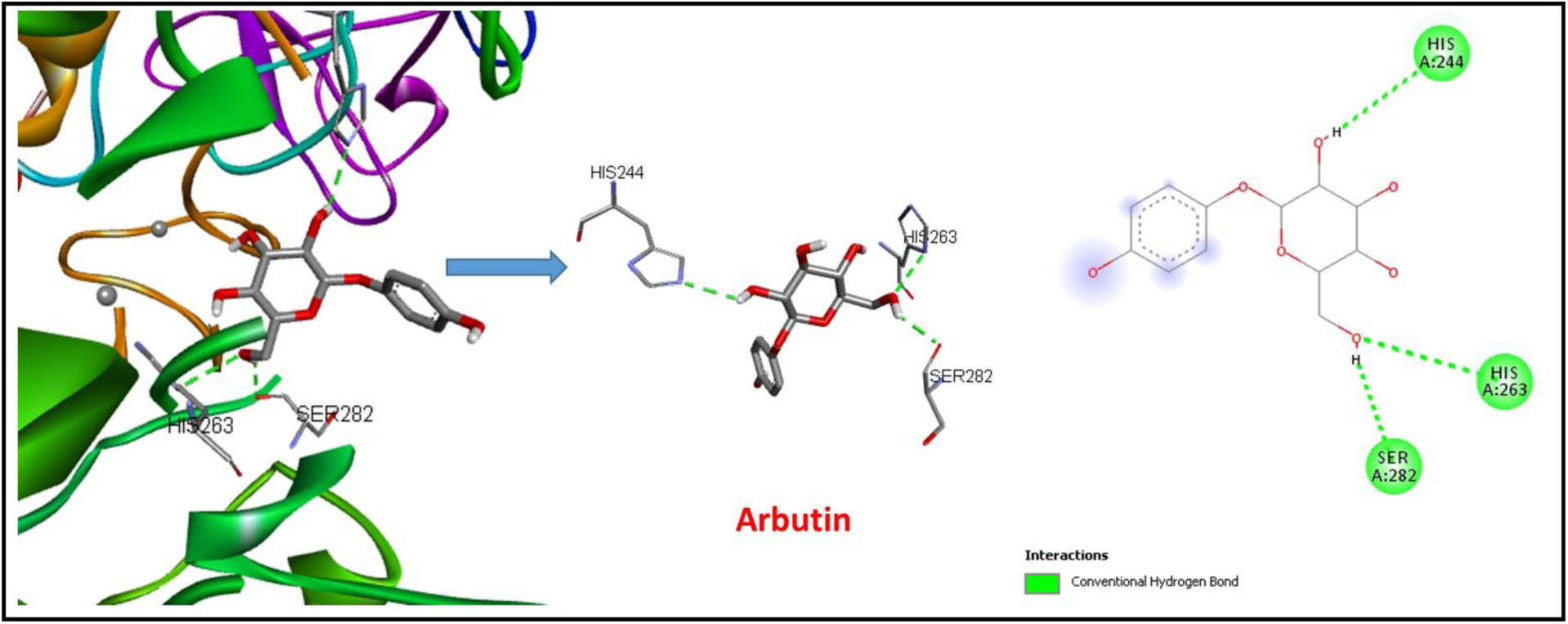
Representation of the putative binding modes of arbutin in the “pocket” of human tyrosinase model. The green dotted lines indicate H-bond interaction and the two gray spheres indicate copper ions at the active site.

**Figure 11:**
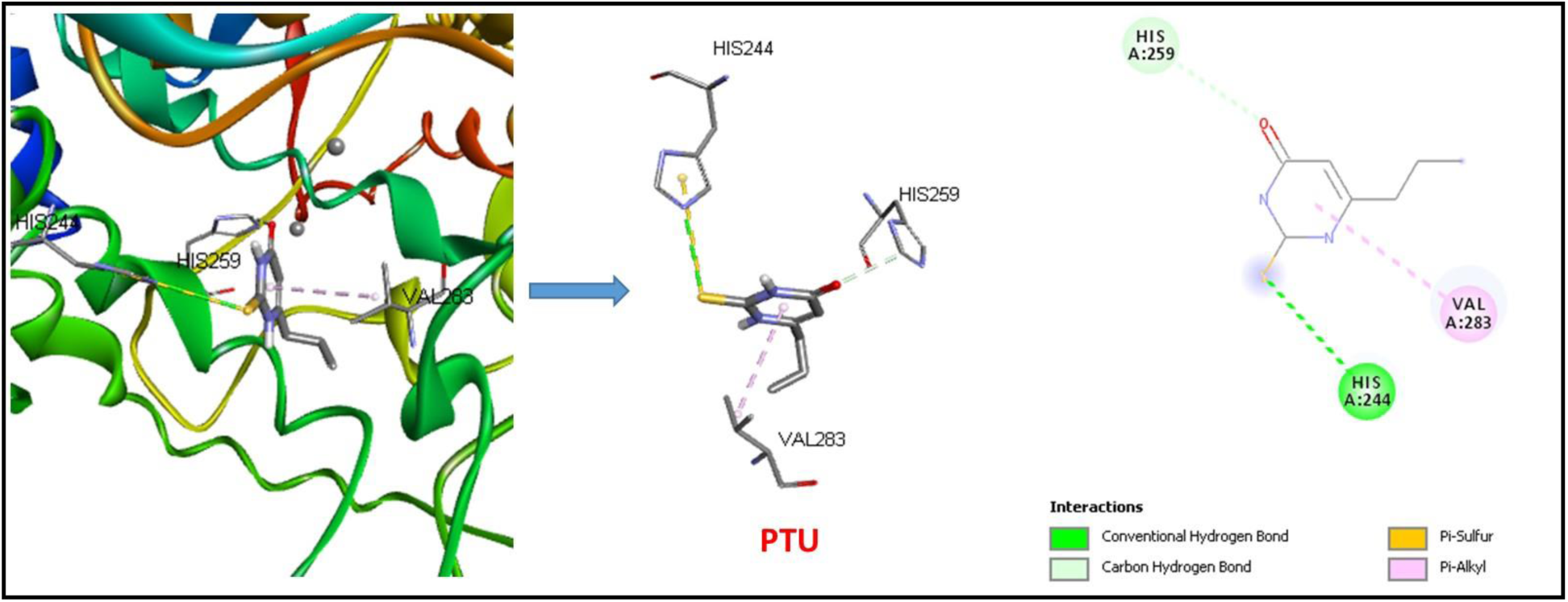
Representation of the putative binding modes of PTU in the “pocket” of human tyrosinase model. The green dotted lines indicate H-bond interaction, the pink dotted lines indicate hydrophobic interactions, yellow dotted lines indicate pi-Sulfur interactions and the two gray spheres indicate copper ions at the active site.

Finally, we show that PTU interacts with residues surrounding the hTYR active site at Gln 286, Gln 376, His 285 and Cys 289 (Figure 7.G, H, I and table 1).

Note that the highest binding energy value corresponded to -6.6 kcal/mol for the Fc-diOH interaction with AbTYR. More interestingly, we found that the highest binding energy value was -8.1 kcal/mol for the Fc-diOH interaction with hTYR (Table 1).

## 4. Discussion

Melanogenesis is responsible for skin color and plays an important role in protecting the skin from sun-related damage. However, the excessive pigmentation can have a significant impact on our appearance. Therefore, tyrosinase inhibitors are frequently used for the treatment of skin hyperpigmentation and as skin witening agents [42]. We have already shown that the Fc-diOH is a competitive inhibitor of Sepia tyrosinase activity [34]. However, further studies are needed to better understand the effect of Fc-diOH on melanogenesis and testing it in models closer to humans.

Here, we examined the effect of Fc-diOH on melanogenesis in the murine melanoma cell line B16F10 and in zebrafish embryos. Then, we used the molecular docking analysis to determine whether Fc-diOH can directly bind to tyrosinase and inhibit its activity with higher affinity.

Inhibition of melanogenesis in B16F10 melanoma cells is associated with a reduced amount of intracellular melanin and tyrosinase activity after treatment with concentrations of Fc-diOH that did not affect cell viability. Fc-diOH reduced intracellular tyrosinase activity (32% at 25 nM) more potently than arbutin (15% at 100 µM) (Figure 3.B).

Furthermore, treatment with 6 µM Fc-diOH induced 50% mortality of zebrafish embryos after 2 days post-fertilization (Figure 4). This concentration is very close to the LC_50_ of tamoxifen in zebrafish, which is equal to 8 µM after 2 days post-fertilization [43]. Fc-diOH induces a depigmenting effect on zebrafish embryos from the concentration 0.1 µM that did not affect its development. We suggest that the decreased melanin content in zebrafish embryos and B16F10 cells treated with Fc-diOH could be due to inhibition of tyrosinase activity and/or the inhibition of expression tyrosinase and other melanogenesis enzymes.

Consistent with the experimental results obtained in *in vivo* and *in vitro* models confirming the inhibitory effect of Fc-diOH on tyrosinase activity, we performed a molecular docking study using AbTYR (*A. bisporus* Tyrosinase) and the model predicted hTYR as a target molecules. Recall that all tyrosinases share a type 3 binuclear copper center within their active sites and that each copper atom is coordinated with three histidine residues. Autodock vina molecular docking analysis shows that the *p*-hydroxyphenyl groups of Fc-diOH interact with the AbTYR active site at histidine residues (His 259 and His 263) which are coordinated with copper atoms, and at of residue Ser 282 via an H-bond interaction. Additionally, the *p*-hydroxyphenyl groups of Fc-diOH interact with the active site of AbTYR at residue Val 283 via two hydrophobic interactions (S. Figure 1 A, B, C and table 1).

The docking study used Molegro Virtual Docker suggested that the phenol groups of Fc-diOH form hydrogen bonds with His 244, Thr 84 and Asn 320 of the *A*.*bisporus* tyrosinase pocket binding site [43]. Additionally, the *p*-hydroxyphenyl groups of Fc-diOH interact with the active site of hTYR tyrosinase at residues His 304, Arg 308 and Ile 368 (Figure 7 A, B, C and table 1). The residue Ile 368 of the active site of hTYR corresponds to the Arg 209 of the active site of bmTYR (S. Figure 3) which play a role in binding orientation of the substrate depending on their flexibility and position [41]. These results suggest that the *p*-hydroxyphenyl groups of Fc-diOH make close contacts with the active site of tyrosinase, which could be due to structural homology between the substrate (L-tyrosine) and the *p*-hydroxyphenyl group of Fc-diOH.

Arbutin, a hydroquinone glucoside compound, has been tested in pharmaceuticals and cosmetics for its ability to prevent overproduction of melanin [44], interacts with the AbTYR active site at Ser 282, as the Fc-diOH interaction, and at the level of residues His 262 and His 244 via the H-bond interaction (S. Figure 1. D, E, F and table 1). Furthermore, the hydroxyphenyl group of arbutin interacts with the hTYR active site at residues His 202, His 367 and Val 377, this result may explain the efficiency of arbutin in inhibiting melanogenesis in B16F10 cells. Additionally, the standard for depigmentation of Zebrafish embryos PTU interacts with the active site of AbTYR at His 259, His 244 and Val 283 (S. Figure 1. G, H, I and table 1). Noting that Fc-diOH interacts with the AbTYR site in the same binding site of arbutin and PTU and it ineracts at the same residue Ser 282 as we found for arbutin interaction and at the same residues His 259 and Val 283 as we found for the PTU interaction. (table 1).

According to the predictive docking energy and binding residues revelation, this supports the conclusion that Fc-diOH exhibits stronger tyrosinase inhibition than arbutin and PTU due to its better binding energy value and its higher number of interactions with amino acid residues of the the active site of hTYR and AbTYR. Therapeutically, Fc-diOH has an anti-proliferative effect on hormone-dependent and hormone-independent breast cancer cells (27), on leukemia cells (45) and on B16F10 melanoma cell lines (38) but it has a depigmenting effect through its potential inhibitory effect on tyrosinase activity (13) and melanogenesis. Fc-diOH may be useful as a potential depigmentation agent for various hyperpigmentation disorders. It is possible because Fc-diOH was found to have low toxicity levels on healthy cells (46, 47).

In conclusion, this study describes for the first time the inhibitory effect of Fc-diOH on melanogenesis in B16F10 cells and in the zebrafish embryo by decreasing melanin synthesis and intracellular tyrosinase activity. Furthermore, molecular docking reveals that the *p*-hydroxyphenyl groups of Fc-diOH make close contacts with the active site of tyrosinase via H-bonds and hydrophobic interactions. This explains the significant inhibitory effect of Fc-diOH on melanogenesis and could be of great interest for a topical formulation in the cosmetic industry for the treatment of various hyperpigmentation disorders.

## CRediT authorship contribution statement

Emna Ketata: Writing – original draft, Methodology, Investigation, Formal analysis, Software, Visualization, Conceptualization, Data curation. Aissette Baanannou and Pascal Pigeon: Methodology, Writing – review & editing, Investigation, Validation. Wajdi Ayadi, Aref Neifar, Siden Top, Saber Masmoudi, Mehdi El Arbi, Gérard Jaouen: Methodology, Writing – review & editing. Ali Gargouri: Writing – review & editing, Supervision, Methodology, Investigation, Conceptualization, Validation, Project administration, Funding acquisition.

## Acknowledgements

We deeply thank Pr. Naziha Marrakchi for the gift of B16F10 melanoma cells. This work has received financial support from “Ministry of Higher Education, Scientific Research and Technology, Tunisia” granted to “Laboratory of Eukaryotic Molecular Biotechnology”, Centre of Biotechnology of Sfax, Tunisia.

